# Secretory trafficking maintains the organelle band for spindle separation during Arabidopsis meiosis

**DOI:** 10.1101/2024.12.03.626687

**Authors:** Ewa Weronika Piskorz, Yingrui Ma, Jun Yi, Maria Cuacos, David Kradolfer, Claudia Köhler, Hua Jiang

**Author notes:** Corresponding authors (Hua Jiang). These authors contributed equally (Ewa Weronika Piskorz, Yingrui Ma).

## Abstract

Ploidy reduction is a critical aspect of meiosis, essential for the successful development of haploid germline cells. Notably, cellular processes during male meiosis vary between the major groups of flowering plants, with eudicots exhibiting simultaneous cytokinesis without an interkinesis phase, unlike monocots. Recent studies on the Arabidopsis *jason* (*jas*) mutant revealed that an organelle band plays a key role in spindle separation during simultaneous cytokinesis, though the mechanism is not fully understood. Here we identify a novel *jas* suppressor mutant, *peleus* (*pele*), which restores haploid pollen production in the *jas* background. PELE encodes a vacuole-localized protein that contributes to spindle separation by balancing the vacuolar pathway and secretory trafficking to the organelle band. JAS, PELE, and the ubiquitin-like protein UBQL coordinate the delivery of proteins and membrane components to the organelle band. Specifically, PELE and UBQL antagonistically fine-tune trafficking to the vacuole, while JAS adjusts the flow of secretory trafficking towards the organelle band. Our findings underscore the critical role of secretory trafficking in maintaining organelle band integrity and in ensuring precise spindle positioning during meiosis II. Our work provides novel insights into the regulation of the organelle band and spindle separation, enhancing our understanding of polyploidy generation in plants.

## Introduction

Sexual reproduction, a fundamental process in plants, is essential for maintaining genetic diversity and facilitating plant adaptation to changing environments (McCormick 2004). Central to this process is meiosis, a specialized type of cell division that produces haploid gametes from diploid parental cells. During meiosis, chromosomes undergo recombination and segregation, ensuring ploidy reduction and the accurate distribution of genetic material to form haploid (1n) gametes. The process of meiosis in monocotyledonous plants typically involves successive cytokinesis, where reductional division is followed by the first cytoplasmic division, resulting in a “transitory dyad” with a cell wall separating the chromosome groups (De Storme and Geelen 2013). Subsequently, equational division occurs within each dyad cell, culminating in the formation of a tetrad of gametes. In dicotyledonous plants like *Arabidopsis thaliana* (Arabidopsis), cytokinesis is simultaneous, with a common cytokinetic event dividing the mother cell into four daughter cells at the end of telophase II (Rodkiewicz and Duda, 1988; Brown and Lemmon, 1996; Zhang et al., 2011). This type of cytokinesis requires organelles and vesicles to create a physical barrier known as the organelle band during meiosis II. The organelle band is crucial for preventing interactions between separated homologs and for ensuring proper segregation of sister chromatids by orienting meiotic spindles perpendicularly (Brownfield et al., 2015; Tchórzewska, 2017).

However, deviations from the standard course of meiosis lead to abnormal meiotic products, including unreduced (2n) gametes. Meiosis, particularly in males, occasionally yields 2n gametes with the same chromosome number as the parent cell, resulting in polyploidy (Brownfield and Köhler, 2011; De Storme and Geelen, 2013). Polyploidy is prevalent in plants and contributes to increased genetic diversity and adaptation. In specific Arabidopsis mutants, such as *jason* (*jas*) and *parallel spindle 1* (*ps1*), large quantities of viable, unreduced male gametes are produced due to the premature loss of the organelle band at prometaphase II or metaphase II, respectively (Brownfield et al., 2015; D’Erfurth et al., 2008; De Storme & Geelen, 2011; Erilova et al., 2009). Although the molecular basis of organelle band disruption is not fully understood, cytological analyses have revealed the presence of two meiotic spindles within the common cytoplasm during meiosis II, allowing unrestricted movement and abnormal orientations (Brownfield et al., 2015). Unreduced gametes play an important role in generating polyploids in nature and represent a promising breeding tool (Ramsey & Schemske, 1998; Adams & Wendel, 2005).

In this study, we explored the mechanism of spindle separation and unreduced gamete formation in Arabidopsis. Our recent study on the *jas* suppressor telamon (*tel*) demonstrated that enlarged meiocytes in *jas tel* mutants restored tetrad formation even in the absence of an organelle band, suggesting a link between cell size and spindle separation (Yi et al., 2023). Here, we analyzed another *jas* suppressor, *peleus* (*pele*), and found that it partially restored the organelle band in *jas* meiocytes, maintaining the distance between chromosome groups at metaphase II in male meiocytes. Our findings further suggest that this restoration in *jas pele* mutants depends on secretory trafficking. We showed that PELE, the ubiquitin-like protein UBQL, and JAS work together to fine-tune the delivery of proteins and membrane components to the organelle band. This regulation occurs through a dual mechanism: (1) PELE and UBQL antagonistically control the secretory trafficking pathway to the vacuole, and (2) the interplay between UBQL-PELE and JAS further modulates the flow of secretory traffic toward either the organelle band or the vacuole. Together, these mechanisms precisely regulate the capacity of secretory trafficking to the organelle band, ensuring proper organelle band maintenance and accurate spindle separation during male meiosis II in Arabidopsis. These results reveal previously unknown roles for secretory trafficking in male meiosis and provide valuable insights into the regulation of the organelle band, spindle separation, and broader mechanisms contributing to polyploidy generation in plants.

## Results

### *pele* suppresses unreduced gamete formation

To investigate the genetic factors influencing unreduced gamete formation in plants, we conducted a forward genetic screen to identify suppressors of the *jas* mutation in Arabidopsis that forms a lot of 2n pollen. In this screen, we identified a mutant named *pele*, which in the *jas* background predominantly produced 1n pollen (Fig. 1A and 1B). This resulted in significantly decreased pollen grain size in *jas pele* compared to *jas* (Kolmogorov–Smirnov test, *P* < 10^−5^) (Fig. 1B). The *jas* mutant exhibited 45% dyads and 33% triads, with all microspores from dyads and one-third from triads developing into 2n pollen. By contrast, the *jas pele* mutant showed significantly reduced numbers of triads and dyads (Chi-square test, *P* < 10^−5^) (Fig. 1C and 1D), indicating that *pele* largely suppresses the male meiotic defect of the *jas* mutant.

**Figure 1.**
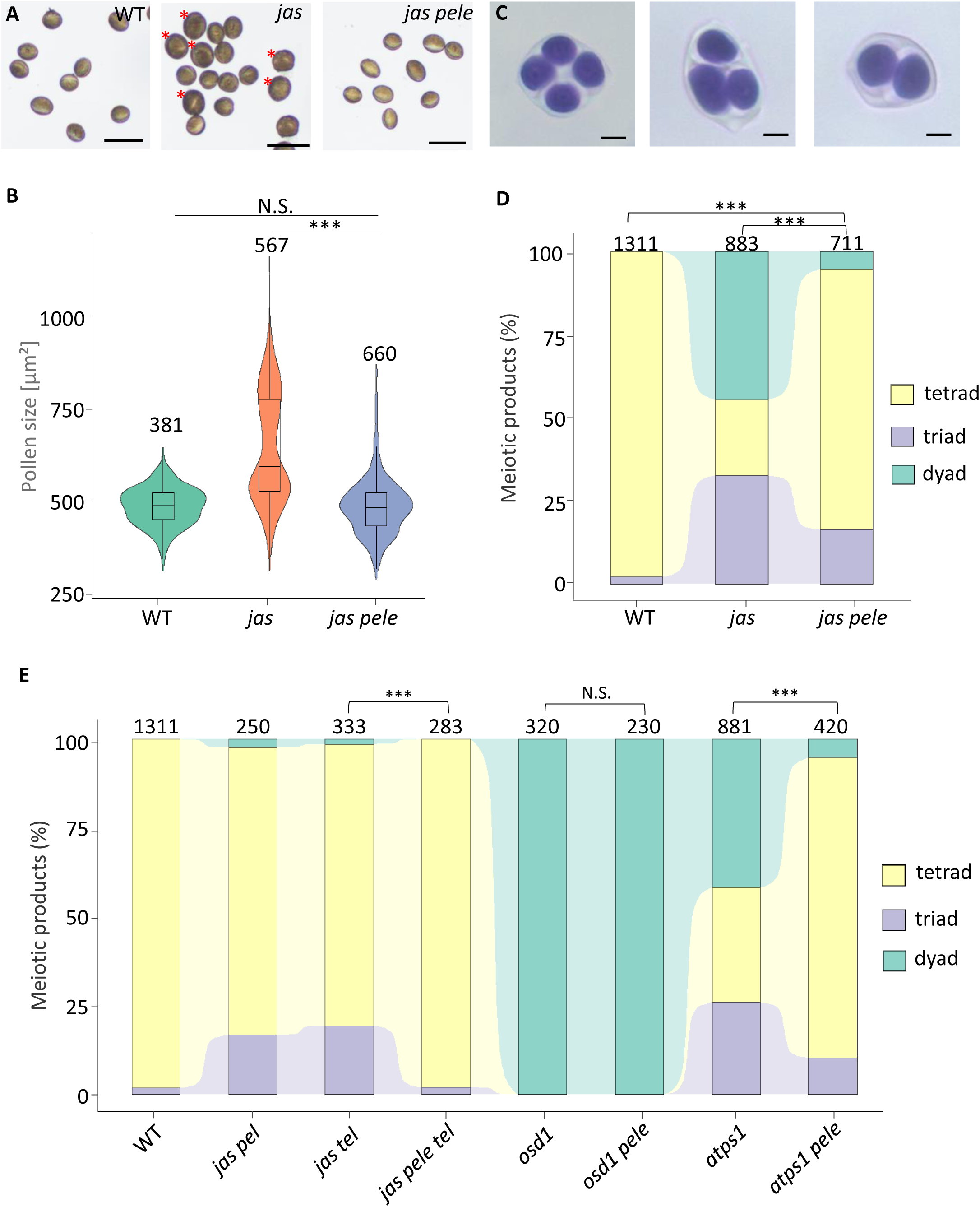
Phenotypic analysis of *jas pele*. (A) Photographs of pollen. Red asterisks indicate diploid pollen. Scale bar, 50 µm. (B) Box plot showing the distribution of pollen size in WT, *jas*, and *jas pele*. The lower and upper hinges of the boxplots correspond to the first and third quartiles of the data, the black lines within the boxes mark the median, whiskers represent Minimum and Maximum, and circles represent outliers. Asterisks mark significance (Kolmogorov–Smirnov test, *P* < 10^-5^). Numbers above indicate the number of meiotic products observed.(C) Photographs of meiotic products stained with toluidine blue. From left to right: tetrad, triad, dyad. Scale bar, 10 µm. Numbers above indicate number of meiotic products observed. (D) Analysis of tetrad formation, with the number of analyzed meiotic products indicated above.E) Tetrad formation of *osd1*, *ps1*, and their progeny from crosses with *pele* and/or *jas*. Significance of the differences was tested by Chi-square test. Asterisks mark significance (*P* < 10^-5^), ns, not significant. Numbers above indicate the number of meiotic products observed.

Recently, we reported that the *tel* mutation suppressed 2n pollen formation within the *jas* background by augmenting the size of meiocytes (Yi et al., 2023). To test if *pele* and *tel* act in the same pathway, we combined them to generate the triple mutant *jas pele tel*. This combination led to a significant increase in the proportion of tetrads compared to the *jas pele* mutant (Chi-square test, *P* < 10^−5^), implying that the mechanism by which *pele* suppresses 2n pollen formation is distinct from that of *tel* (Fig. 1E). Mutations in *OMISSION OF SECOND DIVISION 1* (*OSD1*) also cause unreduced gamete formation in Arabidopsis (d’Erfurth et al., 2008; d’Erfurth et al., 2009) by preventing chromatid separation in meiosis II. We investigated whether *pele* could also suppress the meiotic defect of the *osd1* mutant. The number of dyads and triads in *osd1 pele* double mutants did not substantially differ from those of the single mutants (Fig. 1E), indicating that *PELE* and *OSD1* act in different pathways. By contrast, *pele* significantly suppressed the male meiotic defect of the *ps1* mutant (Chi-square test, *P* < 10^−5^)*. ps1* produced 26% triads and 41.6% dyads, which decreased to 10.5% and 5.2%, respectively, in the *ps1 pele* mutant. Therefore, *pele* suppressed unreduced gamete formation in *jas* and *ps1* mutants, both of which are related to spindle separation during meiosis II.

### *pele* rescues spindle separation in the *jas* mutant

The *jas* mutant has a defect in the separation of chromosome groups and spindles in meiosis II, resulting in the formation of dyads and triads (Brownfield et al., 2015). Given that *pele* suppressed the formation of abnormal meiotic products in the *jas* background, we examined whether the separation of chromosome groups was recovered in *jas pele* by preparing chromosome spreads, which allowed us to visualize chromosomes. The progression of male meiosis I was similar in the wild type (WT), *jas*, and *jas pele* (Fig. 2A, left panel), which is consistent with the finding that *jas* produces unreduced gametes during meiosis II. During male meiosis II in the WT, two chromosome sets aligned at the cell equator, separated by the distinct organelle band. Sister chromatids then segregated into four distal poles of the cell, where they decondensed into haploid nuclei (Fig. 2A, right panel). In the *jas* mutant, the chromosomes became closed or fused, leading to the formation of unreduced meiotic products (Fig. 2A, right panel). By contrast, in *jas pele* during metaphase II, chromosome groups were separated, similar to the WT. Hence, chromosome separation in meiosis II was recovered in *jas pele*.

**Figure 2.**
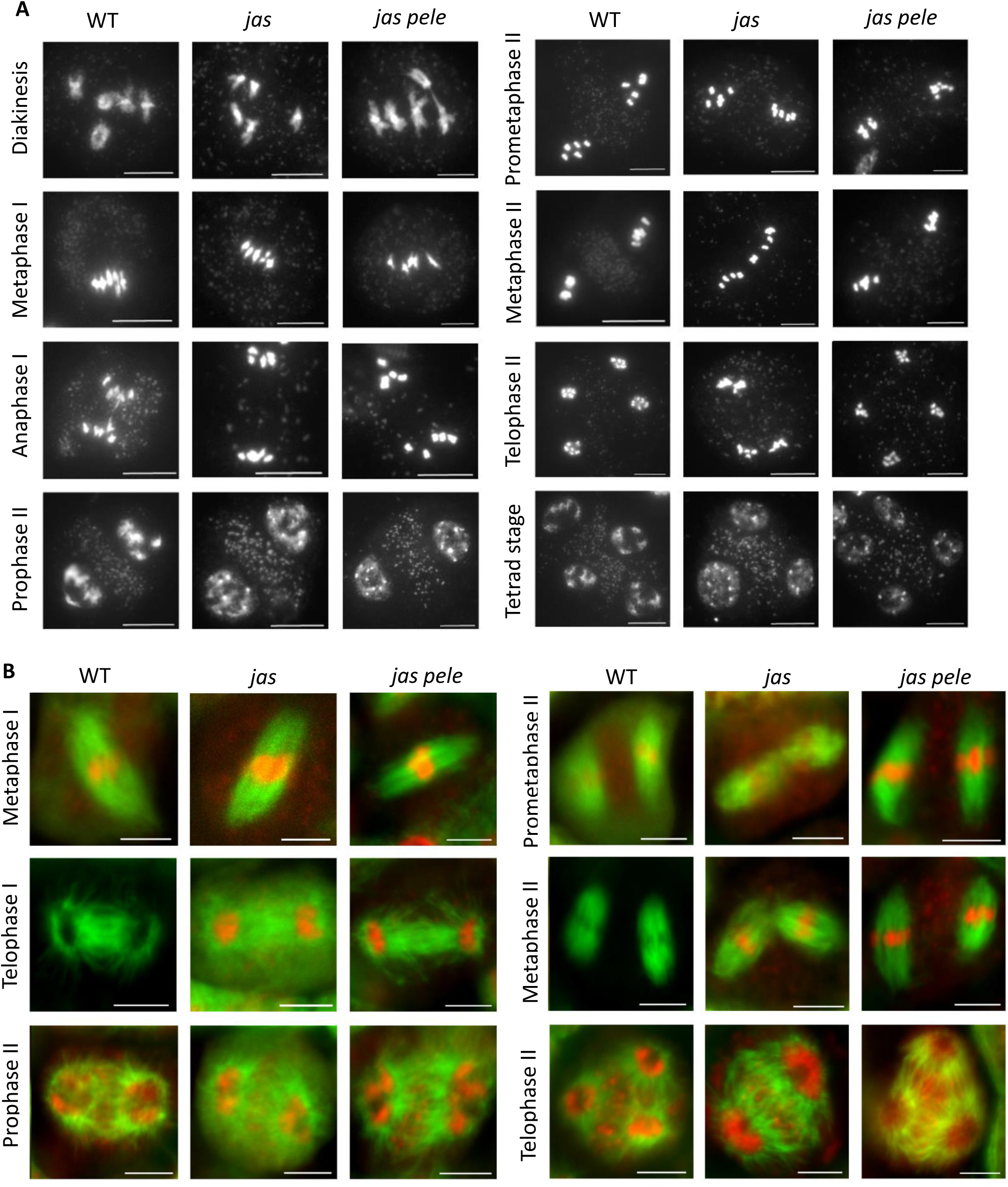
Meiotic progression in *jas pele*. (A) Chromosome spreads of wild-type (WT), *jas,* and *jas pele* meiocytes. Scale bars, 10 µm. (B) Microtubule dynamics during meiosis in WT, *jas*, and *jas pele.* Green signals are from the microtubular cytoskeleton, and red (false color) signals are chromosomes. Very faint or no red signals in some images are due to weak DAPI staining. Scale bars, 5 µm.

The closed or fused chromosome groups in *jas* is associated with abnormal spindle positioning (Brownfield et al., 2015). Therefore, we explored whether *jas pele* also restored spindle positioning. To visualize meiotic spindles and determine their orientation, we used GFP-TUA6 marker lines in the WT, *jas*, and *jas pele* backgrounds. In the WT, meiotic spindles at metaphase II positioned themselves perpendicularly (*n* = 89, Fig. 2B), which led to equal distribution of sister chromatids into four newly forming daughter cells. By contrast, *jas* mutants in addition to normally separated spindles (*n* = 43), displayed tripolar (n = 32) or fused spindles (*n* = 14), giving rise to triads or dyads. However, *jas pele* mostly formed separated, perpendicular spindles, with tripolar spindles constituting only 12.3% of detected conformations (*n* = 14 for a total of *n* = 114; Fig. 2B), and no meiocytes contained fused spindles. We also observed the dynamics of spindle positioning in WT, *jas*, and *jas pele*. In contrast to the stable, separated spindles in WT meiocytes (Movie A), the spindles in *jas* were initially separated but soon became fused, which is in line with the proposed function of JAS in maintaining the organelle band structure (Movie B, C). By contrast, spindle separation was maintained during male meiosis in *jas pele* (Movie D), revealing that *pele* rescued the defects in spindle positioning in the *jas* background.

### *pele* partially rescues the organelle band

Given that JAS functions to separate spindles by maintaining the structure of the organelle band, we investigated whether this maintenance is rescued in *jas pele*. To better visualize the organelle band and further test the effect of *pele*, we stained meiocytes in intact anthers with DAPI to avoid removing the cytoplasm and thereby altering the 3-D structure of meiocytes in chromosome spreads. Although DAPI primarily stains nuclear DNA, it also binds to mitochondrial DNA, meaning the signal in the organelle band may partly reflect mitochondrial structures. Consistent with the chromosome spreads, the DAPI-stained organelle band was well defined in the WT (Fig. 3A), whereas it was partially or often completely absent in *jas*, while it became partially restored in *jas pele* (Fig. 3B and 3C). We counted metaphase II meiocytes containing each type of the organelle band. We considered the organelle band to be “partial” when more organelles were detected between the chromosome sets than in the surrounding cytoplasm. Whereas 100% of WT meiocytes contained complete bands, both *jas* and *jas pele* meiocytes exhibited only a partial or absent organelle band. However, in *jas pele*, we detected partial bands in 58% of the cells (*n* = 82), whereas in *jas*, only a small subset of cells (15%, *n* = 106) contained partial bands (Fig. 3D). The finding that nearly 60% of *jas pele* meiocytes contained a partial organelle band at metaphase II could explain why this mutant produced 82% tetrads, as some meiocytes without an organelle band were still able to undergo correct sister chromatid separation.

**Figure 3.**
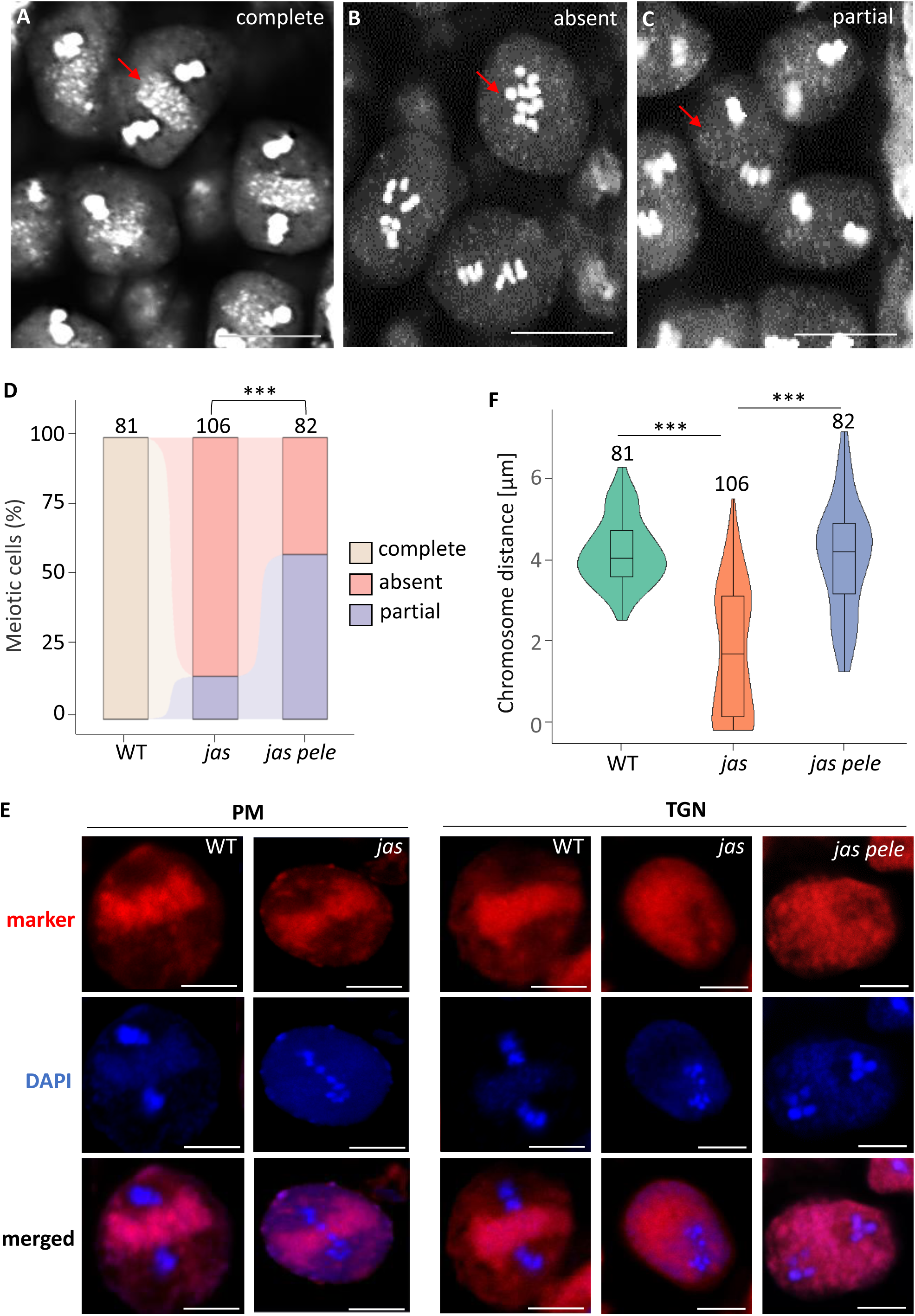
Organelle band types. (A–C) Analysis of whole anthers at metaphase II in WT (A), *jas* (B), and *jas pele* (C), highlighting the three types of organelle bands observed (arrows). Scale bar, 10 µm. (D) Percentage distribution of metaphase II meiocytes with complete, partial, or absent organelle bands observed in WT, *jas*, and *jas pele*. Significance of the differences was tested by Chi-square test. Asterisks mark significance (*P* <10^-5^), ns, not significant. Numbers above indicate the number of meiotic cells observed. (E) Localization of the makers of the plasma membrane (PM), PIP1;4, and trans-Golgi network (TGN), VTI12, in defined genotypes. Scale bar, 10 µm. (F) Box plot showing the distribution of distance between two groups of chromosomes at metaphase II. The lower and upper hinges of the boxplots correspond to the first and third quartiles of the data, the black lines within the boxes mark the median, whiskers represent Minimum and Maximum, and circles represent outliers. Asterisks mark significance (Kolmogorov–Smirnov test, *P* <10^-5^). Numbers above indicate the number of meiotic products observed. (G) Localization of PM and TGN markers in meiocytes at the metaphase II stage in WT, *jas*, and *jas pele*.

In addition to the localization of DAPI-stained organelles, we also observed markers of the plasma membrane (PM) and the *trans*-Golgi network (TGN), which are concentrated in the organelle band (Brownfield, Yi et al. 2015). While chromosome groups are fused in *jas* mutants, the localization of the PM marker in *jas* was similar to wild type (WT) (n = 15 observed meiocytes for jas, n = 8 for WT)) (Fig. 3E, left panel). Thus, disrupted spindle separation in *jas* mutants did not appear to correlate with the localization of the PM marker. In contrast to its distribution in WT (n=23 observed meiocytes), the TGN marker was dispersed in the cytoplasm of *jas* male meiocytes (n=21) (Fig. 4E, right panel), which correlates with the dispersed DAPI-stained organelles and disrupted organelle band. Therefore, we further examined the localization of the TGN marker in *jas pele* mutants. Consistent with the partial recovery of the organelle band, chromosome groups were separated by TGN marker-associated material (n=10 observed meiocytes), although not as densely packed as in WT (Fig. 3E, right panel). Thus, the distribution of the TGN marker was partially restored, coinciding with the partial recovery of the organelle band in *jas pele* mutants.

**Figure 4.**
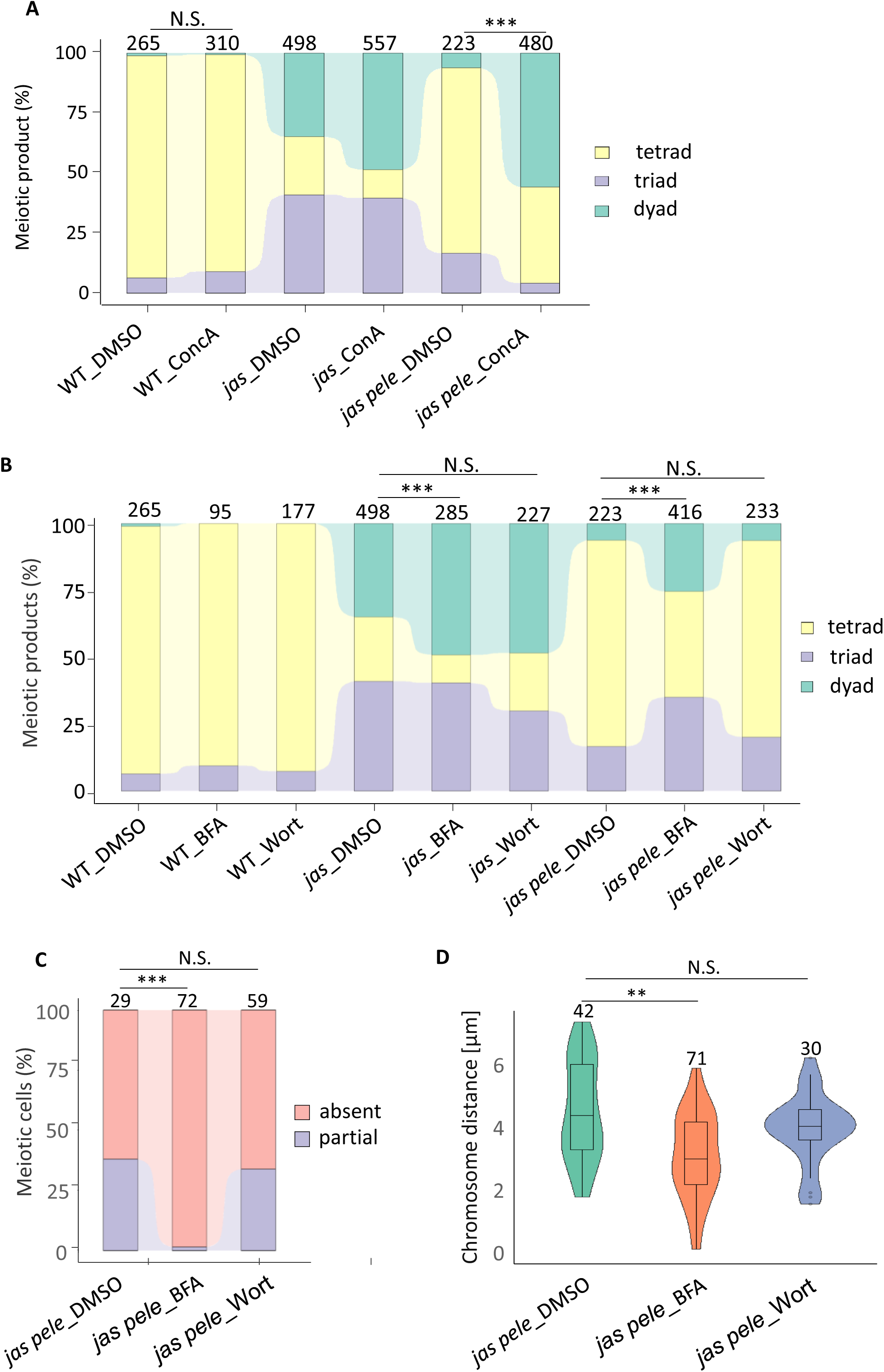
The effect of vesicle trafficking inhibitors on meiosis in the mutants. The effect of vesicle trafficking inhibitors on meiosis in the mutants. (A) Analysis of tetrad formation in WT, *jas*, and *jas pele* after subjecting the inflorescences to Concanamycin A (ConA), and the control DMSO. Numbers above indicate the number of meiotic products observed. (B) Analysis of tetrad formation in WT, *jas*, and *jas pele* after subjecting the inflorescences to Brefeldin A (BFA), and Wortmannin (Wort), and the control DMSO. Numbers above indicate the number of meiotic products observed. (C) Percentage distribution of metaphase II meiocytes with complete, partial, or absent organelle bands observed in *jas pele* after subjecting the inflorescences to vesicle trafficking inhibitors. Significance of the differences was tested by Chi-square test. Asterisks mark significance (*P* <10^-5^), ns, not significant. Numbers above indicate the number of meiotic cells observed. (D) Box plots showing the distribution of distance between two groups of chromosomes at metaphase II in BFA, Wortmannin, and DMSO-treated *jas pele*. The lower and upper hinges of the boxplots correspond to the first and third quartiles of the data, the black lines within the boxes mark the median, whiskers represent Minimum and Maximum, and circles represent outliers. Asterisks mark significance (Kolmogorov–Smirnov test, *P*< 10^-5^). Numbers above indicate the number of meiotic cells observed.

The partial recovery of the organelle band was confirmed by measuring the distance between metaphase II chromosome sets. In *jas* mutants, chromosome sets were frequently either closer than in WT or even touching (n = 106 observed meiocytes, Fig. 3F). In contrast, chromosome sets in *jas pele* mutants were consistently located at a distance of at least 1.5 μm, significantly longer than in *jas* mutants (Kolmogorov–Smirnov test, *P* < 10^−5^) (Fig. 3F). Thus, the *pele* mutation suppressed the *jas* phenotype by partially restoring organelle positioning and maintaining the separation of chromosome sets at metaphase II.

### Vesicle trafficking maintains the organelle band in male meiocytes

The organelle band consists of multiple types of organelles and vesicles, which are typically associated with vesicle trafficking. Therefore, we investigated whether the restoration of the organelle band in *jas pele* mutants depends on vesicle trafficking. If this is the case, inhibiting the relevant trafficking pathways should diminish the rescue of the organelle band and tetrad formation defect in *jas pele* mutants. To test this hypothesis, we examined the effects of vesicle trafficking inhibitors on meiotic outcomes in *jas pele* mutants by treating inflorescences with inhibitor solutions and assessing the meiotic products thereafter. Initially, we utilized Concanamycin A, a chemical that impacts TGN function by blocking all TGN-associated pathways, including the endocytic and the secretory trafficking pathways (Dettmer et al., 2006; Reichardt et al., 2007). This treatment notably weakened the suppressive effect of *pele* on the *jas* mutant phenotype (Chi-square test, *P* < 10^−5^), leading to a significant increase in the percentage of dyads and triads from ∼30% (treated with DMSO) to ∼70% (Chi-square test, *P* < 10^−5^) (Fig. 4A). This suggests that *pele* indeed restores spindle separation through its effect on vesicle trafficking.

To further investigate which specific pathway *pele* affects, we treated inflorescences with Wortmannin and Brefeldin A (BFA), which inhibit the endocytic pathway and the secretory or recycling pathways, respectively. While Wortmannin did not affect the phenotype of *jas pele* mutants (Fig. 4A), BFA significantly increased the percentage of triads and dyads (to ∼60%) (Chi-square test, *P* < 10^−5^). Furthermore, to verify the influence of the inhibitor treatment on organelle band recovery, we assessed the percentage of the partial organelle band. Consistent with the tetrad formation results, BFA treatment suppressed the partial organelle band in *jas pele* mutants (Chi-square test, *P* < 10^−5^), whereas Wortmannin did not (Fig. 4B). Consequently, BFA, but not Wortmannin, significantly reduced the distance between two groups of chromosomes at metaphase II (Kolmogorov-Smirnov, *P* < 0.01) (Fig. 4C).

While treatment with Concanamycin A and BFA notably induced the production of abnormal meiotic products in *jas pele* mutants, these chemicals had no effect on male meiosis in WT plants, suggesting that vesicle trafficking partially contributes to maintain the organelle band in WT plants and that another mechanism independently stabilizes the organelle band, unrelated to vesicle trafficking.

### *PELE* encodes a vacuole-localized protein

To identify *PELE*, we map-based cloned *pele* and identified a C-to-T substitution in the 18th exon of the gene AT5G11700. This substitution caused a tryptophan to be replaced by a stop codon. Derived cleaved amplified polymorphic sequence (dCAPS) analysis and Sanger sequencing performed on *jas* and *jas pele* DNA confirmed that the substituted nucleotide originated from ethyl methanesulfonate (EMS) treatment (Fig. S1A and B).

To further test whether AT5G11700 is the *PELE* gene, we crossed the at5g11700 T-DNA insertional mutant with *jas* and examined the double mutant progeny in the F_2_ generation. The double mutants exhibited a phenotype similar to *jas pele*, producing approximately 80% tetrads (Fig. S1C). To verify this, we generated a construct containing the WT genomic AT5G11700 sequence along with its 1.5-kb promoter and used this construct to transform *jas pele* plants. AT5G11700 complemented the *jas pele* phenotypes in the T_1_ and T_2_ generations, restoring 1n pollen formation and increasing the number of dyads (Table S1). Thus, we identified *PELE* as AT5G11700.

To elucidate the role of PELE in meiosis, we crossed *jas pele* with WT and obtained the *pele* single mutant. Analysis of pollen size indicated that *pele* produced uniformly haploid pollen (Fig. S2A), which developed from the reduced meiotic products (97.8% tetrads, *n* = 458, Fig. S2B). Moreover, we performed chromosome spreads of the single mutant. At metaphase II, two chromosome sets were separated by a complete organelle band characteristic of the WT (Fig. S2C), and no other meiotic abnormalities were detected. Hence, the *pele* mutation affects meiotic progression only in the presence of the *jas* or *ps1* mutation.

PELE consists of 1,453 amino acids and lacks known domains that could suggest its biological function. Online sources, such as SMART (embl-heidelberg.de), predicted the presence of a signal peptide at its N terminus and at least four transmembrane domains (Fig. S3A), indicating that it is a membrane-localized protein. Previous studies localized PELE to the vacuole membrane and annotated it as a putative multi-transmembrane tonoplast protein of unknown function (Endler et al., 2006; Shimaoka et al., 2004). To examine the *PELE* expression pattern, we fused the *PELE* promoter and genomic sequence with the *β-GLUCURONIDASE* (*GUS*) gene, transformed this construct into WT plants, and subjected the progeny of the resulting transformants to GUS staining. The *PELE* promoter was nearly constitutively expressed in all plant organs, including roots (Fig. S3B, C), rosette leaves (Fig. S3D), cauline leaves (Fig. S3E), and inflorescences (Fig. S3F), throughout all stages of plant development.

We further observed PELE-GFP localization in male meiocytes by fusing the *PELE* promoter and genomic sequence with GFP. PELE-GFP was dispersed in the cytoplasm of male meiocytes (Fig. 5A), partially overlapped with the area of the organelle band, suggesting the functional relationship between PELE and the regulation of organelle band. In contrast to the pattern in male meiocytes, in roots, we observed co-localization of PELE-GFP with markers for the late endosome/prevacuolar compartment and the tonoplast but not with other markers (Fig. 5B, Fig. S4, Table S2). Moreover, PELE-GFP formed vesicle-like structures that emitted bright florescence that, however, did not co-localize with any of the Wave line markers for the ER, Golgi, TGN/early endosomal compartments or PM (Fig. S4), indicating that PELE-GFP was localized at other types of organelles or aggregated in cytoplasm. In summary, PELE localizes in the late endosome and tonoplast in root cells, consistent with its functional role in vesicle trafficking during male meiosis.

**Figure 5.**
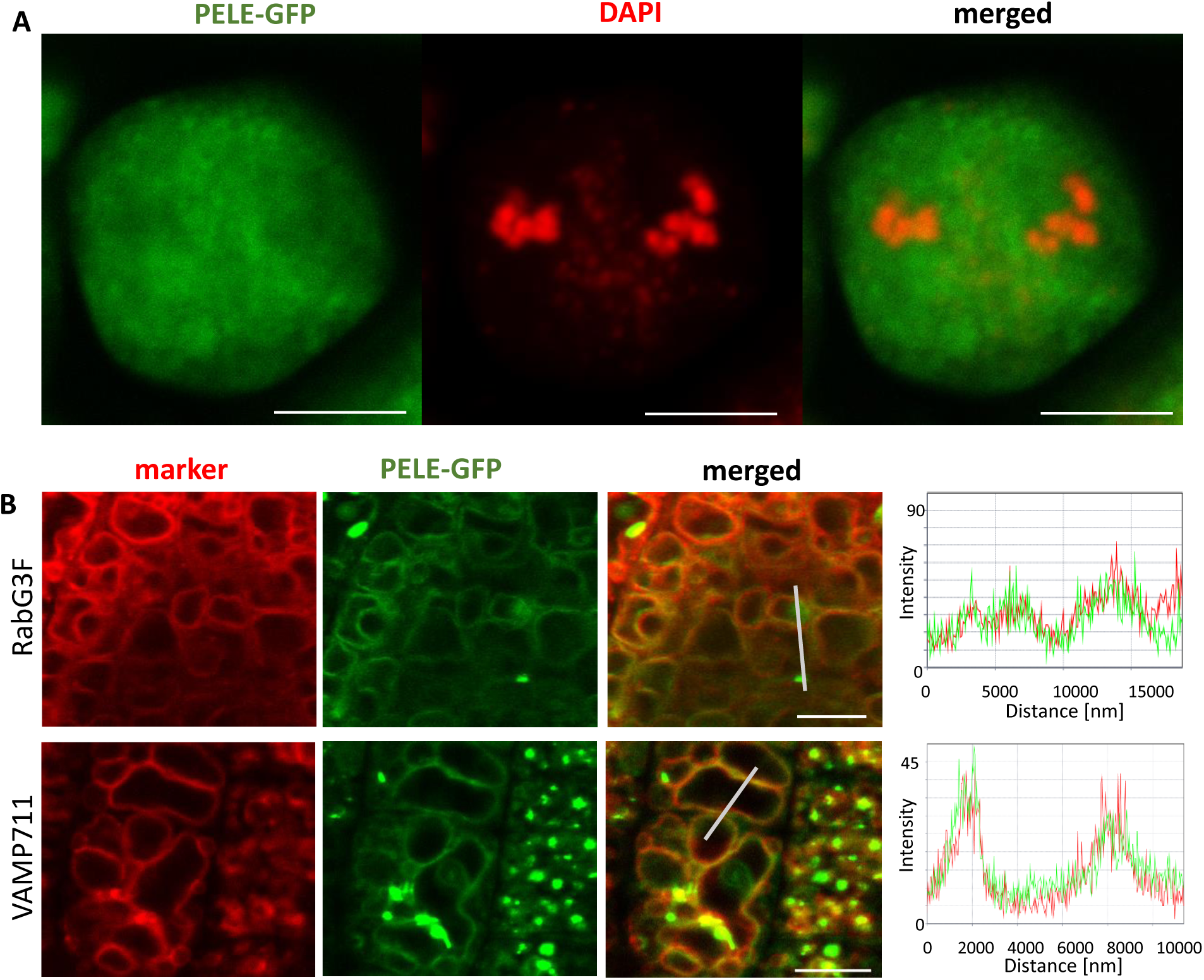
PELE localization in meiocytes and roots. (A) Localization of PELE-GFP at metaphase II in WT meiocytes. Green signals are from PELE-GFP, red signals are from chromosomes. Scale bar, 10 μm. (B) Localization of PELE-GFP in roots. RabG3F, late endosome and tonoplast; VAMP711, tonoplast. Scale bar, 5 μm.

### UBQL together with PELE fine-tune secretory trafficking to the organelle band

Pharmacological and genetic analyses suggest that secretory pathways are crucial for regulating the organelle band. The tonoplast localization of PELE-GFP indicates its role in this process. Since the *pele* mutation rescues defects in secretory trafficking, we propose that PELE helps balance vacuolar and secretory pathways. This mutation may reroute cargoes that would normally go to the vacuole toward the organelle band, potentially explaining the partial rescue of the *jas* phenotype. To understand how PELE influences this process, we investigated potential PELE interactors. Through the BioGRID protein interaction database (Oughtred et al., 2019), we identified UBQL (AT5G42220), a ubiquitin-like protein, as a potential candidate. Consistent with this, AlphaFold2 predictions indicated an interaction between PELE and UBQL, specifically through amino acids W^1028^, R^1029^, and G^1031^ of PELE, and E^758^, Q^753^, and G^766^ of UBQL (Fig. 6A and 6B). We validated this interaction through a split-ubiquitin yeast two-hybrid assay. Since full-length PELE inhibited yeast growth, we tested a fragment of PELE (PELE^909-1296^), which includes the predicted interaction sites. In addition to testing UBQL and PELE^909-1296^, we included UBQL with an empty vector as a negative control and UBQL with Nub1 as a positive control. The yeast containing UBQL and PELE^909-1296^ showed growth similar to the positive control, with noticeably more colonies compared to the negative control. Thus, our findings support an interaction between UBQL and PELE in yeast consistent with an interaction predicted by *in silico* analyses.

**Figure 6.**
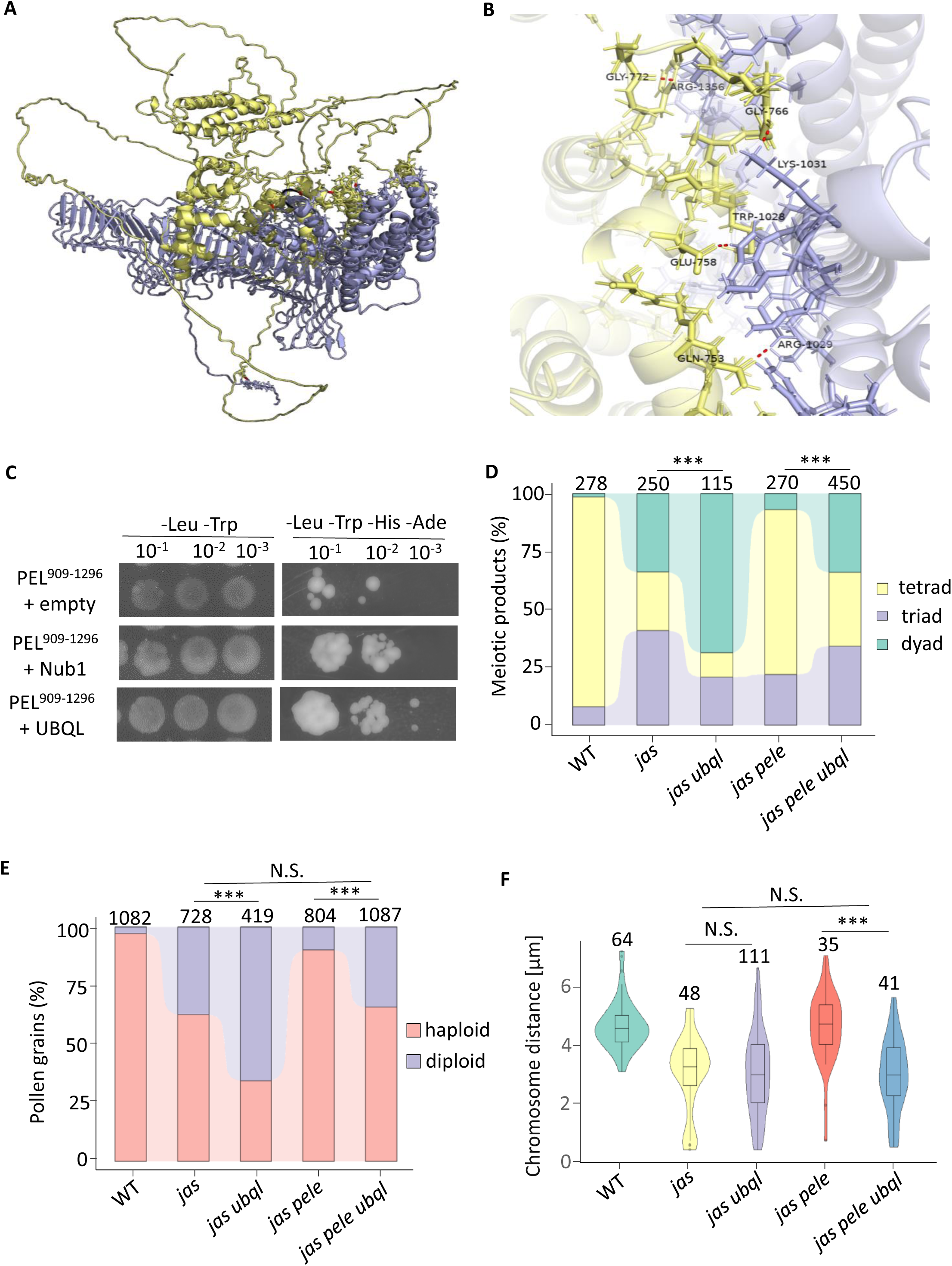
Function of UBQL in organelle band regulation. (A) Predicted protein interaction between PELE and UBQL using AlphaFold2. UBQL is shown in yellow, and PELE is shown in purple. Magnified image of the protein interaction site in (A). Polar bonds are indicated by red dotted lines. (C) Yeast two-hybrid assay results showing interaction between UBQL and the PELE (909-1296) fragment. (D) Analysis of tetrad formation in WT, *jas*, *jas ubql*, *jas pele*, and *jas pele ubql* mutants. Numbers above indicate the number of meiotic products observed. (E) Predicted percentage distribution of haploid and diploid pollen based on the tetrad analysis results in (D). Statistical significance was tested using the Chi-square test. Asterisks indicate significant differences (P < 10⁻⁵); ns, not significant. Numbers above indicate the number of meiotic products observed. (F) Violin plots showing the distribution of the distance between two chromosome groups at metaphase II in WT, *jas*, *jas ubql*, *jas pele*, and *jas pele ubql*. The lower and upper hinges of the boxplots correspond to the first and third quartiles of the data. The black lines within the boxes represent the median, whiskers represent the minimum and maximum values, and circles represent outliers. Asterisks indicate significant differences (Kolmogorov–Smirnov test, P < 10⁻⁵). Numbers above indicate the number of meiotic cells observed.

To further investigate the role of UBQL in secretory trafficking-mediated spindle separation, we crossed *jas pele* with *ubql* to generate *jas ubql* and *jas pele ubql* mutants. Despite the interaction between UBQL and PELE, the *ubql* mutation significantly increased the production of dyads under the *jas* background (Chi-square test, P < 10⁻⁵) (Fig. 6D). This suggests that UBQL positively affects secretory trafficking, in contrast to PELE, which may negatively affect the process potentially due to a primary role in the vacuolar pathway. Given these opposing roles, we hypothesized that mutating both PELE and UBQL in the *jas* background would neutralize their individual effects, leading to levels of dyads and triads in the *jas pele ubql* mutant similar to in the *jas* mutant. As expected, the percentage of dyads in *jas pele ubql was* significantly lower than in *jas pele*, making it closer to the levels observed in *jas* mutants (Fig. 6D). Each triad produces one diploid (2n) pollen and two haploid (1n) pollen, while each dyad generates two 2n pollen. Based on these patterns, we speculated on the proportions of 1n and 2n pollen from the tetrad analysis (Fig. 6E). As anticipated, the percentage of diploid pollen changed significantly when *PELE* and *UBQL* were mutated individually in the *jas* background (Chi-square test, P < 10⁻⁵). However, no significant difference was found between *jas* and *jas pele ubql*, suggesting a balancing effect when both *PELE* and *UBQL* are disrupted as both the vacuolar and the secretory pathway may be affected by the respective mutants. In line with the tetrad and pollen analysis results, chromosome distance at metaphase II was comparable between *jas* and *jas pele ubql*, whereas the distance was shorter in *jas pele ubql* than in *jas pele* (Fig. 5F). These findings strongly support the conclusion that UBQL and PELE antagonistically fine-tune secretory trafficking to the organelle band, where UBQL takes over a positive function in contrast to PELE that negatively affects the process.

## Discussion

Our findings highlight the function of secretory trafficking in maintaining the integrity of the organelle band during male meiosis in Arabidopsis. Previous data revealed localization of JAS to the organelle band and its association with various organelles in root cells, suggesting that JAS is involved in organelle band maintenance (Brownfield et al., 2015; Cabout et al., 2017). Here, we identified PELE as factor negatively impacting organelle band formation or maintenance. We show that mutations of *PELE* restore the organelle band in the *jas* mutant background. This restored organelle band in *jas pele* is disrupted by inhibiting secretory trafficking. PELE is associated with the tonoplast in root cells, consistent with a possible function in vacuolar vesicle trafficking. Unlike the significant effects on organelle band formation observed in the *jas* and restored in the *jas pele* mutant, none of the chemical treatments with vesicle trafficking inhibitors on their own disrupted organelle band formation and spindle separation in WT plants. This may suggest the existence of another mechanism independent of vesicle trafficking that stabilizes the organelle band. Considering the substantial disruption of the organelle band in the *jas* mutant, we propose that JAS is responsible for both, secretory trafficking-dependent and independent mechanisms. Therefore, in the *jas pele* mutant, restored secretory trafficking resulted in a partial organelle band, while inhibition of vesicle trafficking suppressed the organelle band. In future studies, exploring the nature of the secretory trafficking-independent mechanism in organelle band formation promises to yield intriguing insights.

Secretory trafficking typically delivers proteins to the plasma membrane or vacuole (Zhang et al. 2019). However, during cytokinesis, it is redirected towards the zone where the future cell plate will form (Kanazawa and Ueda 2017). The enrichment of the plasma membrane marker PIP1;4 in the organelle band at metaphase II (Brownfield et al., 2015) suggests a similar redirection of vesicle trafficking to the central zone of male meiocytes in Arabidopsis. Notably, treatment with BFA, an inhibitor of ER-to-Golgi transport, suppressed the rescued spindle separation observed in *jas pele* mutants, indicating that the secretory pathway, including ER-to-Golgi transport, is involved in organelle band regulation. Interestingly, the effect of BFA treatment in male meiocytes differed from its effect in seedlings, where the functional redundancy between the BFA-sensitive GNOM and the BFA-resistant GNL1 limits the observable impact of BFA (Van Damme et al., 2008). In male meiocytes, however, the effect of BFA was already evident in the wild-type background, suggesting that GNL1 does not compensate for GNOM in ER-to-Golgi transport during male meiosis, unlike in seedlings.

Wortmannin treatment did not produce a significant effect on *jas pele*, suggesting a minor role of endocytic transport in organelle band regulation. The localization of the endomembrane marker PIP1;4, which is likely transported via endocytic transport, remains unaffected when the organelle band is disrupted in *jas*, supporting the notion that the import of endomembrane-localized proteins alone is insufficient to maintain the organelle band structure. However, we cannot entirely rule out the involvement of the endocytic pathway in organelle band regulation, as Wortmannin does not completely inhibit endocytosis, leaving open the possibility that endocytic vesicles contribute to this process (Leshem et al. 2007). Therefore, the relative contributions between secretory and endocytic trafficking to organelle band regulation remain to be further determined.

The transport of newly synthesized proteins and membrane components to the plasma membrane or vacuole is tightly regulated during plant development (Zhang et al., 2019). PELE is primarily localized in late endosomes and the tonoplast in root cells, suggesting a similar role in male meiocytes, although the vacuole morphology is altered. During somatic cytokinesis, vacuoles undergo substantial changes in organization, size, and volume, with an 80% reduction in vacuole volume compared to prometaphase cells (Segui-Simarro and Staehelin, 2006). Similar dynamics may occur in male meiocytes, potentially altering the localization and function of PELE compared to root cells. At metaphase II, as a tonoplast-localized protein, PELE is likely primarily involved in transport to or at the tomoplast, while JAS is required for secretory trafficking toward the organelle band. To modulate the activity of PELE, UBQL antagonizes PELE by blocking secretory transport to the vacuole. On the other hand, *pele* mutations may reroute transport of some cargoes to the vacuole into the secretory pathway contributing to partial restoration of the organelle band. Potentially antagonistic roles of PELE and UBQL thus may help fine-tune the capacity of secretory trafficking to the vacuole (Fig. 7) and to the PM/organelle band, balancing the two pathways. In *jas male* meiocytes, excessive vacuolar trafficking disrupts the organelle band, but in *jas pele* mutants, UBQL partially rescues band function by restoring some trafficking to the organelle band. In *jas ubql* mutants, the absence of UBQL enhances the ability of PELE to reroute trafficking to the vacuole, worsening organelle band disorganization and affecting spindle separation. This secretory trafficking is fine-tuned by two mechanisms: PELE and UBQL balance vacuolar transport, and their interplay with JAS regulates trafficking to the organelle band, ensuring proper spindle separation during male meiosis II in Arabidopsis.

**Figure 7.**
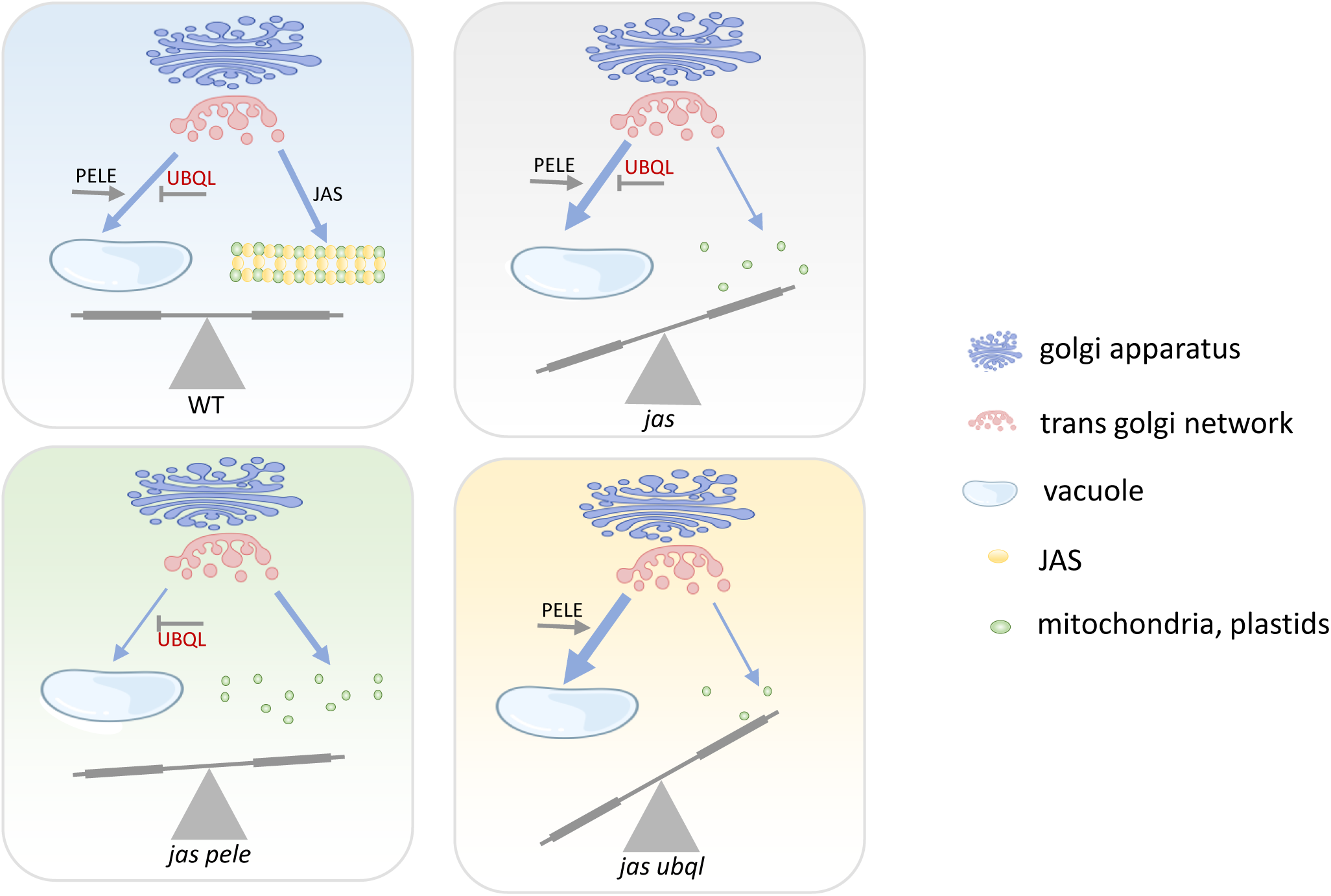
Model of secretory trafficking in organelle band organization. The figure illustrates how the balance between PELE-UBQL and JAS regulates the transport of proteins and membrane components to both the organelle band and the vacuole. PELE and UBQL exhibit an antagonistic interaction that refines the trafficking pathway towards the vacuole. Disruption in JAS, PELE, or UBQL causes an imbalance in this regulatory system, leading to altered secretory trafficking and affecting the organization of the organelle band.

The resemblance of the organelle band to the cytokinesis-derived cell plate formed during successive cytokinesis raises questions about the connection between organelle band formation and cytokinesis. Whereas vesicle trafficking, particularly exocytosis, is essential for cell plate formation during cytokinesis (Reichardt et al., 2007), successful cell plate formation does not occur in male meiocytes with simultaneous cytokinesis (De Storme and Geelen, 2013). To ensure proper spindle separation during male meiosis II, plants may have evolved unique mechanisms involving vesicle trafficking to maintain organelles in a band between spindles, a process likely dependent on JAS. The specificity of JAS to eudicots (Cabout et al., 2017), with monocots using a different mechanism during meiosis II, highlights the diverse regulatory pathways involved in plant reproduction. Together, our results generate novel insight into the role of secretory trafficking in maintaining the organelle band, allowing proper spindle separation during simultaneous cytokinesis.

## Materials and Methods

### Plant materials and growth conditions

Seeds of the Arabidopsis (*Arabidopsis thaliana*) T-DNA insertional mutants *peleus* (SALK_209914C), the *jas-3* allele (SAIL_813_H03), and the *ps1-1* allele (SALK_078818) were obtained from the Nottingham Arabidopsis Stock Centre (NASC). The marker lines of organelles from the Wave line collection harboring endomembrane system markers fused with mCherry (Geldner et al., 2009) were obtained from NASC. The WT Arabidopsis ecotype Col-0 was employed.

Seeds were stratified for 48 hours at 4°C in water in the dark, sown in soil, and propagated in a phytotron under a 16-h-light/8-h-dark photoperiod. The temperature was set to 22°C in the daytime and 19°C at night, the light intensity was 115 μmol·m^−2^·s^−1^, and the relative humidity was 70%.

EMS mutagenesis of *jas-3* seeds, genetic mapping, sequencing, and mutant identification were carried out as previously described (Kradolfer et al., 2013; Yi et al., 2023).

### Pollen phenotyping

To measure pollen size, a drop of 1× PBS with 0.1% Triton X-100 was placed on a microscope slide and two to three open flowers were dipped into the solution to release pollen. After covering the sample with a coverslip, the pollen was imaged at 10× magnification under a SMZ1500 Nikon optical microscope. The size of each pollen grain was measured in ImageJ using a self-made algorithm, in which the ‘Area’ category reflects the 2-D area of a pollen grain.

### Cytological analysis

To count the meiotic products, whole inflorescences were fixed in Carnoy’s fixative (ethanol:acetic acid 3:1) for at least 3 h at room temperature. After two washes with 1× PBS, two to three of the largest white buds were dissected under a stereoscope, and the anthers were cut into pieces using sharp needles or a razorblade. The meiotic products were stained with 0.1–0.2% toluidine blue solution, viewed at 10× magnification under a stereoscope, and manually counted. Chromosome spreads were prepared by fixing the inflorescences in Carnoy’s fixative. Two to three ca. 0.5 mm long white buds were separated from the rest of the buds and digested for 20 min at 37°C in 10 mM sodium citrate buffer, pH 4.5, with 0.3% cellulase and 0.3% pectinase. The buds were washed tree times in 1× PBS and placed on a slide under a stereoscope. The meiotic anthers were released and macerated with sharp tweezers. The homogenous suspension of cells was quickly incubated in 60% acetic acid on a hot plate set to 45°C, gently spread on a slide, and washed with a drop of fresh, ice-cold Carnoy’s fixative. The slide was left to dry and covered with 2 µm/mL DAPI (Roth) in antifade solution. Meiocytes were examined using a Zeiss Axioplan 2 fluorescence microscope equipped with a Plan-Apochromat 63x/1.40 oil immersion objective.

Whole meiotic anthers were analyzed by fixing inflorescences undergoing meiosis in Carnoy’s fixative as previously described and removing intact anthers from the meiotic buds. The anthers were incubated in DAPI buffer (0.4 µL of 2 µg/mL DAPI in 1× PBS + 0.15% Triton X-100) for at least 20 min at room temperature in the dark. The samples were sealed under a coverslip and imaged using a Zeiss LSM780 confocal microscope (Carl Zeiss, Jena, Germany) with a C-Apochromat 40x/NA 1.2 water immersion objective and DAPI filter.

To visualize various cellular components fused with fluorescent tags, inflorescences undergoing meiosis were fixed in 100% methanol at –20°C for 20 min. The samples were washed with ice-cold 1× PBS and kept on ice until processing. Intact meiotic anthers were prepared for analysis using the ‘whole anther analysis’ procedure, with occasional dissecting using a razorblade or gentle squashing with a coverslip to enhance weak signals. Imaging was performed using a Zeiss LSM780 confocal microscope equipped with a C-Apochromat 40x/NA 1.2 water immersion objective. The DAPI filter was used for imaging chromosomes and the organelle band, eGFP for meiotic spindles and PELE-GFP, and mCherry for endomembrane system marker lines.

### Inflorescence culture and inhibitor treatment

The main stems of 6-week-old Arabidopsis inflorescences were cultured on medium comprising 200 mL of 1× MS salts, 3% sucrose (w/v), and 1× vitamins (Feldmann, 1991). The pH was adjusted to 6.0 with KOH, and the medium was autoclaved at 121°C for 15 minutes prior to use. Inhibitors were prepared as a master mix. Concanamycin A (ChemCruz) powder was dissolved in DMSO to create a 200 μM stock solution. Wortmannin (Abcam) powder was dissolved in DMSO to generate a 3 mM stock solution. Both solutions were stored at –20°C and fully dissolved before the experiment. Brefeldin A solution 1000× (BioLegend) was freshly diluted in 9× DMSO to create the master mix prior to use. The treatment medium was prepared by adding master mix to the inflorescence culture medium at a ratio of 1:100, resulting in a final concentration of 2 μM Concanamycin A, 30 μM Wortmannin, and 7 μM Brefeldin A. The prepared treatment medium was aliquoted into 0.5-mL Eppendorf tubes and sealed with a sheet of adhesive foil to prevent contamination. A needle was used to create a hole in the middle of each tube. Inflorescences in excellent condition were selected, carefully cut with a blade, and immediately inserted into the hole in each 0.5-mL Eppendorf tube for treatment. The culture system was incubated under standard conditions for 22 h. After this incubation period, the treated inflorescences were fixed and analyzed. The fixed samples were stored at a maximum of 4°C for up to 1 week.

### Lightsheet Fluorescence Microscopy

Lightsheet Flurescence Microscopy (LSFM) was performed using a Lightsheet 7 (Carl Zeiss GmbH) equipped with two pco.edge 4.2 sCMOS cameras (PCO AG) according to previous reports (Ovecka et al. 2015, Valuchova et al. 2020, Feng et al. 2023) with minor modifications. ∼0.5 mm Arabidopsis flower buds were carefully detached from the inflorescence, dissected to remove outer sepals, and inserted into 1.5 mm (inner diameter) glass capillaries (size 3, green mark; Carl Zeiss GmbH, 701908) containing 1% low melting agarose (Sigma-Aldrich, A9045) prepared in distilled water. Once fitted in the microscope, the agarose cylinder containing the sample was extruded into the imaging chamber filled with distilled water. Temperature in the chamber was set to 21 °C, and no artificial light source was supplemented. Imaging was done with 20x (W Plan-Apochromat 20x/1.0) detection and 10x (LSFM 10x/0.2 foc) illumination objectives and 2.0x zoom. Excitation was done with 0.8% 488 nm laser line. Dual-side illumination was chosen, and pivot scan was turned on to reduce shadowing. Z-stacks covering the whole anther were taken every 45 sec. Images were acquired with ZEN Black 3.1, and processing (selection of single z-stack, drift correction where needed, ROI selection, movie export) was done with ZEN Blue 3.4 (both from Carl Zeiss GmbH).

### DNA library preparation

The leaves of 3-week-old seedlings were snap-frozen and stored in –80°C until DNA extraction. The leaves were pooled and homogenized using a mortar and pestle. DNA was extracted from the sample using a DNeasy Plant Mini Kit from Qiagen according to the manufacturer’s protocol. Four micrograms of high-quality DNA was sonicated with a Bioruptor (Diagenode) into 200- to 300-bp fragments, and the DNA library was prepared using a VAHTS Universal DNA Library Prep Kit for Illumina from Vazyme according to the manufacturer’s protocol. The concentration of the libraries was measured on a Qubit Fluorometer (Thermo Fisher Scientific), and library quality was measured on an Agilent 2100 Bioanalyzer (Agilent Technologies). The libraries were sequenced at Novogene, UK, and at least 15 million reads were produced for each sample (*jas pele* and *jas* as a control). Reads were mapped to TAIR10 Col-0 genomes with HISAT2 (Kim et al., 2019).

### Plasmid construction and genotyping

To generate the genomic *PELEUS* (*PELE*) construct for complementation analysis, a 10,590-bp fragment of AT5G11700, including 1,531 bp of the upstream region, the stop codon, and 570 bp of the downstream region, was amplified from Col-0 genomic DNA with Phanta Max Super-Fidelity DNA Polymerase from Vazyme (forward primer 5’-ATAAGCTTGATATCGAATTCttggaatccgcgaaaatgttatg-3’ and reverse primer 5’-ACTAGTGGATCCCCCGGGgaaggggaagatgagggaatga-3’). The TSK108 entry vector was linearized with *Eco*RI and *Sma*I (ThermoFisher), and the g*PELE* fragment was inserted into the vector using a ClonExpress Ultra One Step Cloning Kit (Vazyme). To generate the gPELE-eGFP and gPELE-GUS constructs, the same genomic locus, but without the stop codon and the downstream region, was amplified and cloned as described (reverse primer 5’-ACTAGTGGATCCCCCGGGCGACTGCCAAAACAGCTCAT-3’). The assembled sequences were transferred into Gateway-compatible destination vectors using LR Clonase (ThermoFisher): PELE into pGWB510, PELE-GFP into pH7FWG;0, and PELE-GUS into pGWB533. After verifying the constructs by Sanger sequencing, Arabidopsis plants were transformed with the vectors using the floral dip method (Bent & Clough, 1998) mediated by *Agrobacterium tumefaciens* strain GV3101. Transgenic lines were selected on ½ MS plates supplemented with 35 μg/mL hygromycin.

To genotype the mutants and transformants, DNA was extracted from 3-week-old plants using the Edwards method (200 mM Tris-HCl pH 7.5, 250 mM NaCl, 25 mM EDTA, and 0.5% SDS) (Edwards et al., 1991) with some modifications, and PCR was carried out with 2× Taq Master Mix from Vazyme. Primers used for genotyping are listed in Table S2.

### GUS staining

WT Col-0 plants were transformed with pGWB533-PELE-GUS, and the resulting seeds were selected on ½ MS plates supplemented with 35 μg/mL hygromycin before being transferred to soil at approximately 2 weeks after germination. Various organs of the positive transformants were collected at different developmental stages, stained with GUS staining solution (50 mM phosphate buffer, 10 mM EDTA, 0.1% Triton X-100, 1 mM potassium ferricyanide III, 1 mM potassium ferrocyanide II, and 1 mg/mL X-Gluc), incubated overnight at 37°C in the dark, and washed with 70% ethanol; WT Col-0 was used as a negative control.

## Acknowledgements

We gratefully acknowledge Amanda Souza Camara for her expert assistance with the protein interaction analysis using AlphaFold2, and Markus Grebe for his valuable feedback that helped improve the manuscript. This research was supported by intramural funding from Leibniz Institute of Plant Genetics and Crop Plant Research (IPK), Gatersleben (to HJ), and a grant from the German Research Foundation (DFG) to HJ (JI 347/5-1).

## Author contributions

E.P., Y.M., and H.J. conceived and designed the experiments, and wrote the manuscript. J.Y. and M.C. executed the experimental procedures. D.K. and C.K. contributed materials. All authors discussed the results and commented on the manuscript.

## Supplemental figure legends

**Figure S1.** Confirmation of the point mutation within AT5G11700. (A) dCAPS analysis of the point mutations in *jas-3* and *jas-3 pele*. (B) PCR product sequencing confirming the point mutation in AT5G11700, which results in the generation of a stop codon. (C) Analysis of tetrad formation in six representative *jas-3* at5g11700 double mutants. The number of analyzed meiotic products is indicated above the bars.

**Figure S2.** Phenotypic analysis of the *pele* single mutant. (A) Distribution of pollen size in the *pele* single mutant and three control lines. (B) Percentage of meiotic products in *pele* and the control lines. (C) Stages of meiosis in *pele*; from left to right: diakinesis, metaphase I, anaphase I, prophase II, metaphase II, anaphase II, telophase II, and the four nuclei stage. Scale bar is 10 μm, which is identical to all photos.

**Figure S3.** Structure of PELE and histochemical GUS staining in tissues of Pro:PELE-GUS plants. (A) Schematic representation of the PELE protein structure predicted by SMART (embl-heidelberg.de). The diagram includes the T-DNA insertion site and most commonly predicted features, such as the four transmembrane domains and the cleavable signal peptide at the N terminus. The amino acid with a point mutation in the *pele* mutant is indicated by a lightning mark. (B–G) Histochemical GUS staining in tissues of Pro:PELE-GUS plants. (B) 7-day-old seedling. (C) 7-day-old seedling, with more visible staining in the root. (D) 14-day-old seedling. (E) Top of a leaf of a 4-week-old plant. (F) Cauline leaf of a 5-week-old plant. (G) Inflorescence of a 5-week-old plant. Scale bars, 1 mm.

**Figure S4.** PELE localization with organelle markers in roots. NIP1, ER and the plasma membrane; Got1p homolog, the Golgi apparatus; VTI12, the trans-Golgi-network and early endosome; RabF2a, late endosome and pre-vacuolar compartment. Scale bar, 5 μm.

**Table S1** Complementation analysis for *jas pele*.

**Table S2** Primers used in the manuscript.

